# Pathogenesis and immune responses to Eurasian avian like H1N1 and influenza D virus in pigs

**DOI:** 10.64898/2026.05.05.722908

**Authors:** Ashutosh Vats, Liu Yang, Ehsan Sedaghat-Rostami, Catherine Hatton, Emily Briggs, Graham Freimanis, Tim Downing, Kristien van Reeth, Basudev Paudyal, Francisco J. Salguero, Wilhelm Gerner, Elma Tchilian

## Abstract

Eurasian avian like H1N1 (EAavH1N1) and influenza D viruses (IDV) with their ongoing evolution and zoonotic potential are a serious threat to animal and human health. Using experimental infection of pigs, we characterized and compared their pathogenesis, and immune responses. EAavH1N1 induced rapid viral clearance, early immune activation, including robust systemic and mucosal antibody responses and increased IFNγ and TNF production. This heightened immune response was associated with more severe pathology of the upper and lower respiratory tract. In contrast, IDV infection resulted in prolonged viral shedding and higher viral titres, with delayed and attenuated cellular immune responses. Single cell transcriptomic analysis of lung further indicated early and persistent suppression of antiviral and innate immune pathways during IDV infection. These findings demonstrate that EAavH1N1 and IDV exhibit distinct viral kinetics, immune activation profiles, and lung responses, providing insight into differences in transmission dynamics, disease severity, and immune control among influenza virus types in swine.

## Introduction

Influenza viruses are a major threat to animal and human health, causing significant morbidity, mortality and economic losses. Their ongoing antigenic and genomic evolution reduces the long-term effectiveness of vaccines and antiviral drugs [1]. Continued circulation and reassortment among humans, livestock, and wildlife drive the emergence of novel zoonotic variants with pandemic potential [2]. Pigs are natural host and a potential mixing vessel at the human - livestock interface. Their respiratory tract expresses α2,3-linked and α2,6-linked sialic acid receptors, as well as 9-O-acetylated sialic acids, making them susceptible to avian adapted and human-adapted influenza A viruses (IAV) and supporting infection by influenza D virus (IDV) [3, 4].

The Eurasian avian-like H1N1 (EAavH1N1) lineage remains a priority among swine IAVs due to its enzootic spread in Europe and Asia and its ability to reassort with other swine influenza viruses [5, 6]. Reassortment with internal genes from the 2009 pdmH1N1 and North American triple-reassortant swine viruses produced genotype 4 (G4) EAavH1N1 viruses, that enable efficient replication in mammalian airways and greater affinity for human-type (α2,6-linked) receptors [7]. These features can facilitate cross-species transmission [7–9]. Surveillance continues to detect sporadic human infections with EAavH1N1-related viruses, including cases without direct pig contact, indicating that spillover can occur through unrecognized modes of transmission [8, 10, 11]. Experimental infection with EAavH1N1 indicates that it can cause respiratory disease in pigs ranging from moderate influenza-like illness with the G1 genotype to more severe lower respiratory tract pathology caused by the G4 genotype [12]. Similarly, studies in mice have shown that recent strains of the G4 EAavH1N1 lineage exhibit higher replication levels in lungs compared with earlier G1 viruses [13].

Influenza D virus has emerged more recently as an important pathogen at the livestock interface. In contrast to influenza A viruses, IDV has a seven-segment genome and a haemagglutinin esterase fusion glycoprotein, HEF, that combines receptor binding, receptor destroying and fusion functions[14–17]. HEF targets 9-O-acetylated sialic acids, which are abundant in cattle and present in pigs [6, 18, 19]. IDV is actively diversifying, with multiple genetic and antigenic lineages circulating and evidence of reassortment [15]. Cattle are the main reservoir, but pigs are susceptible with serological and virological data indicating repeated exposure in domestic and wild swine populations [18, 20–23].) Serological studies show high exposure among cattle- and swine-associated workers, even without sustained human-to-human spread [24–26]. In cattle, IDV pathogenesis has been studied in much greater detail and is generally characterised by mild respiratory disease, efficient transmission and measurable airway innate responses, although some strains can also involve the lower respiratory tract and contribute to more severe disease in the context of coinfection with Mycoplasma bovis [23, 27, 28]. By contrast, in pigs IDV infection has usually been described as subclinical or mild, with replication largely restricted to the upper respiratory tract [18, 20, 21], however recent evidence suggests that some strains may replicate efficiently and transmit between pigs [23, 29, 30].

Although both EAavH1N1 and IDV circulate in swine and are relevant to animal and public health, most studies have focused on virology, transmission, or serology, leaving limited understanding of the immune mechanisms that control infection or drive tissue damage. Here, we used experimental infection with EAavH1N1 and IDV in pigs to define and directly compare their pathogenesis and immune control. We characterised viral replication kinetics, respiratory tract pathology, and local and systemic immune responses, together with lung transcriptomic changes across early and later stages of infection. By analysing these parameters side by side in the natural host, we identified lineage specific differences in lung pathology and immune regulation, providing a more integrated understanding of influenza virus infection in swine.

## Results

### EAavH1N1 causes earlier lung pathology, whereas IDV shows prolonged replication

Fifteen pigs were infected intranasally either with D/swine/Netherlands/PS-497/2021 Influenza D virus (IDV) or A/swine/Gent/274/2020 (EAavH1N1). Six animals were left untreated as controls (**Fig. 1A**). At 1-, 5-, and 12-days post infection (DPI), 5 animals per infected group and 2 controls were euthanised to assess early (1 and 5 DPI) and late events (12 DPI) following infection **(Fig. 1A**). Viral shedding was quantified by plaque assay and qRT–PCR of nasal swabs. Both viruses showed productive infection by 1 DPI, but IDV reached higher titres and persisted until 5 DPI, while EAavH1N1 shedding ended by 3 DPI (**Fig. 1B, S1 B and C**). The nasal shedding of IDV was significantly higher than EAvH1N as analysed by the area under the curve p = <0.0001. No infectious virus or viral RNA were detected by 10 DPI.

**Fig. 1:**
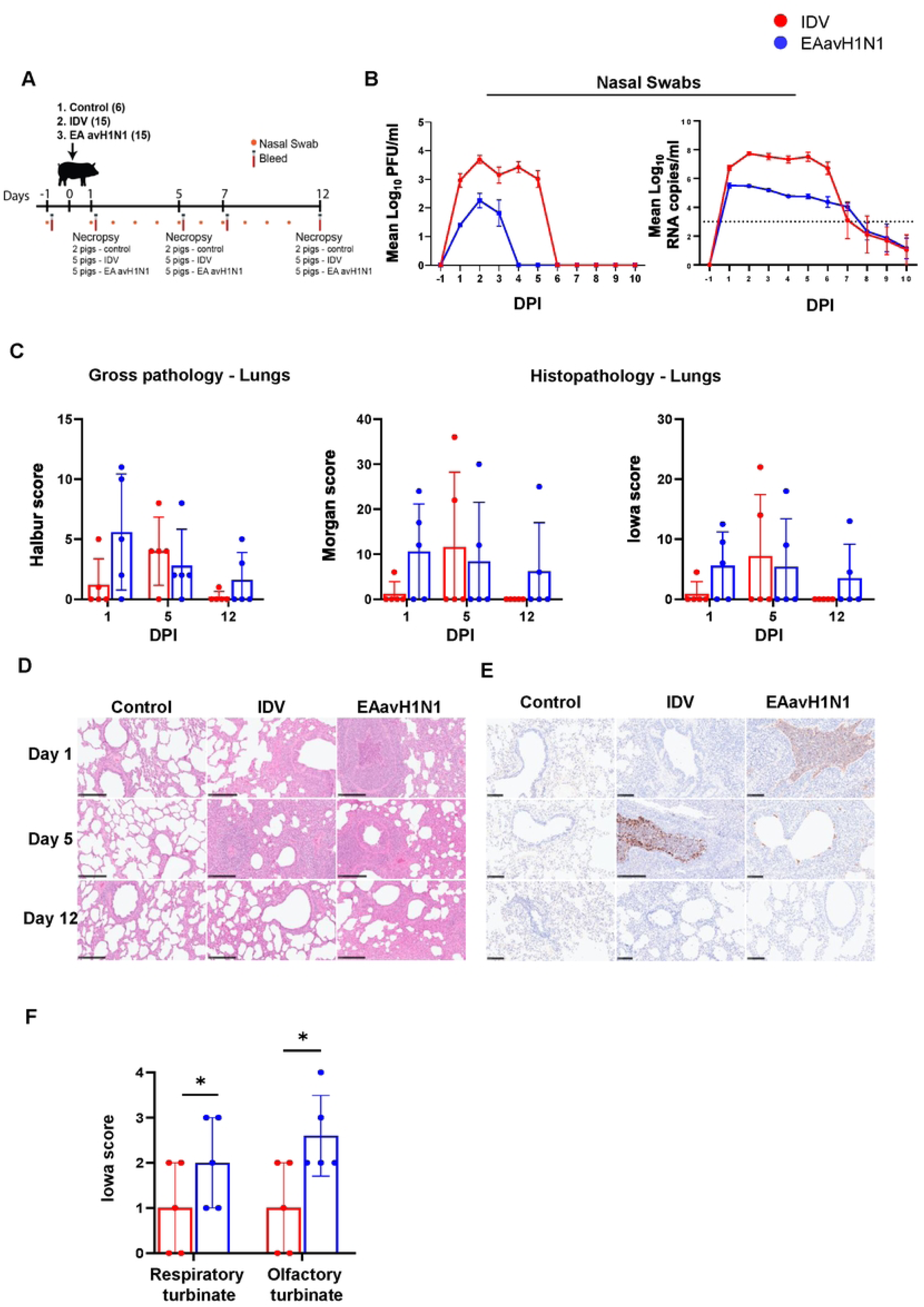
Experiment design, Lung pathology and Viral shedding post IDV and EAavH1N1 infection. (A) Thirty-six pigs were divided into three groups: untreated controls (n=6), avH1N1-infected (n=15), and IDV-infected (n=15). Daily nasal swabs were taken through Day 10 to monitor viral shedding. Necropsies were performed on Days 1, 5, and 12, with 2 control, 5 avH1N1, and 5 IDV pigs per timepoint for lung pathology. (B) Viral titers were measured by plaque assay and RT-qPCR. Swab samples were collected from Day -1 to Day 10 post-infection for both IDV and EAavH1N1 groups. Results are shown as mean Log_10_ PFU/ml or Log_10_ RNA copies/ml. Data are presented as mean ± SEM, with each symbol representing an individual animal. Lung sections from each group were assessed for gross pathology, histopathology (H&E staining), and viral nucleoprotein (NP) distribution by IHC. (C) Gross pathology, and histopathology (Morgan and Iowa). (D) Histopathology revealed bronchiolo-interstitial pneumonia with epithelial necrosis and inflammatory infiltrates. (E) NP-IHC staining observed mainly in the bronchiolar epithelial cells and inflammatory cells infiltrates and exudates (brown staining) (F). Histopathology and immunohistochemistry scores of the respiratory turbinate, and olfactory turbinate in IDV, and EAavH1N1 at 5 DPI. Asterisks indicate significance between groups (*p < 0.05), analyzed by Two-way ANOVA with Bonferroni’s multiple comparisons test (F) for normally distributed data. Scale bar= 100 µm.

Lung pathology was assessed by gross and histopathological scoring as previously described, including immunohistochemistry (IHC) for detection of viral nucleoprotein (NP) [31]. EAavH1N1 induced earlier and more severe pathology, detectable at 1 DPI, peaking at 5 DPI, with milder lesions still present at 12 DPI, while IDV showed milder lesion at 1 DPI, similar lesion at 5 DPI and clearance at 12 DPI **(Fig. 1C and S1A**). Histopathological evaluation showed multifocal broncho-interstitial pneumonia after infection with either virus (**Fig. 1D**). Viral NP was observed within the airway epithelia and within the inflammatory cell infiltration in the alveoli luminae and interalveolar septa, at 1 and 5 DPI in EAavH1N1 and only at 5DPI in IDV infected animals (**Fig. 1E**). EAavH1N1 induced significantly more severe pathology in the upper respiratory tract, particularly in the respiratory and olfactory turbinates, than IDV at 5 DPI (**Fig. 1F**).

Together, these data show that EAavH1N1 induced earlier and more severe pathology in the respiratory tract, while IDV resulted in higher and more prolonged nasal shedding.

### Antibody responses to IDV and EAavH1N1

Virus specific systemic and mucosal IgG and IgA responses were analysed in serum, bronchoalveolar lavage (BAL) and nasal swabs by ELISA and virus neutralization assays **(Fig. 2)**. EAavH1N1 specific IgG in serum was detectable by 5 DPI and peaked at 12 DPI, whereas IDV specific IgG appeared at 7 DPI and peaked at 12 DPI (**Fig. 2A**). IgA titres followed a similar pattern to IgG although at lower titers. Virus specific IgA appeared earlier after EAavH1N1 infection but was higher at 12 DPI in IDV infected animals **(Fig. 2D)**. In BAL, both IgG and IgA were detectable at 12 DPI in IDV infected pigs and were higher compared to EAavH1N1 **(Figs. 2B and E)**. BAL titers were low, reflecting the very diluted nature of sampling. Nasal swabs titers were also low, but detectable at higher levels in IDV samples at 10DPI **(Figs. 2C and F)**. Serum neutralizing antibody activity mirrored the ELISA titers with higher levels in IDV than EAavH1N1 by 12 DPI **(Fig. 2G)**. No neutralisation was detected in the BAL and nasal swabs.

**Fig. 2:**
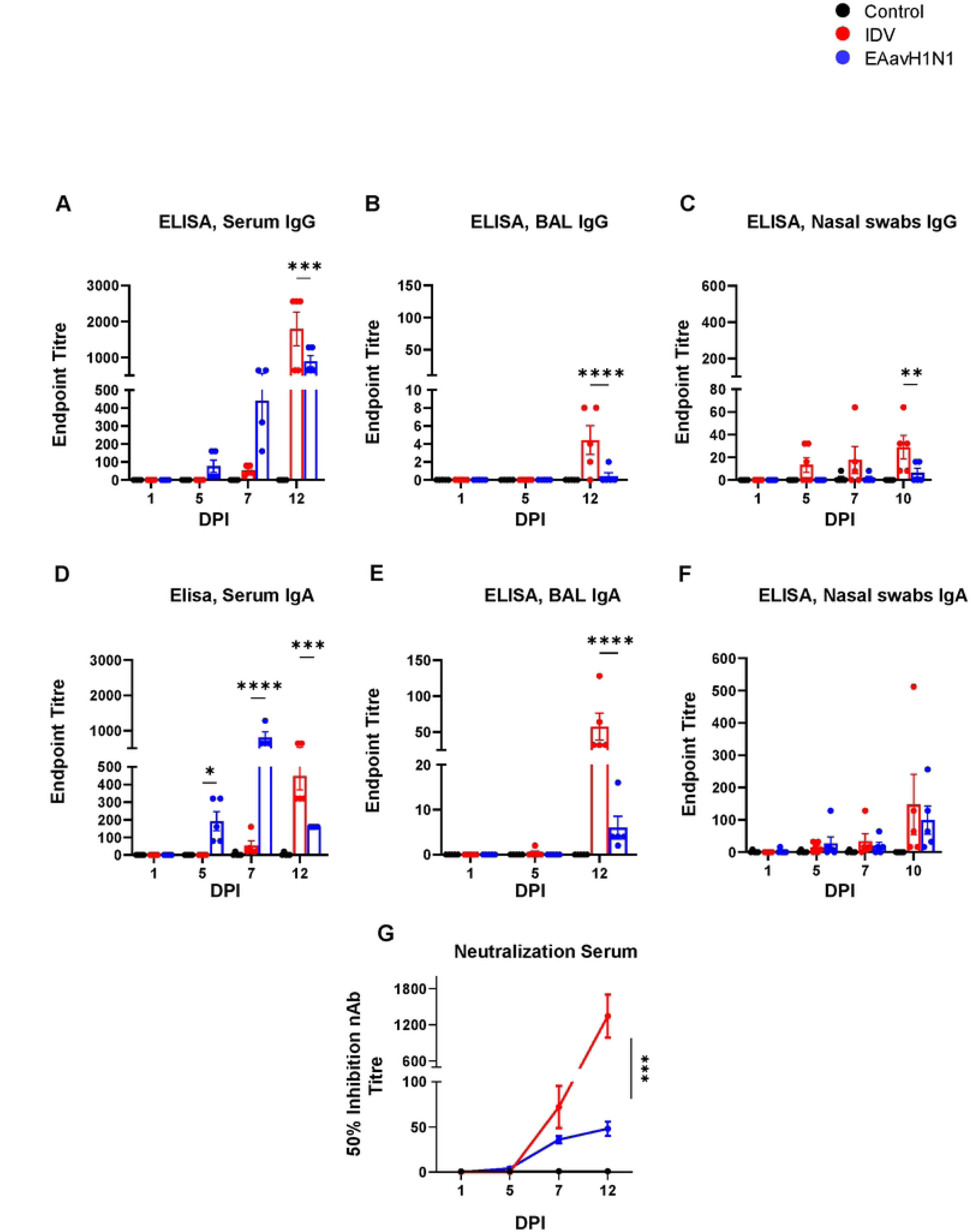
Quantification of IDV and EAavH1N1 -specific antibody responses: lgG and lgA responses were quantified in serum (A, D) , bronchoalveolar lavage (B, E), and nasal secretions (C, F) from pigs at indicated time points after infection with either IDV or EAavH1N1. Antibody levels were measured using virus-specific ELISAs. Data are shown as end-point titres. mean ± SD (n = 5 per group) with symbols indicating individual animal. (G) Neutralizing antibody titter in serum at indicated time points after infection with either IDV or EAavH1N1. Statistical comparisons were performed using *t:wo-way* ANOVA with Bonferroni’s multiple comparisons test for normally distributed data. Asterisks indicate statistically significant differences bet:ween IDV and EAavH1N1 responses at corresponding time points (…p < 0.05, ·••p < 0.01, ·•-p <0.001).

Taken together, these data indicate that EAavH1N1 induced earlier and stronger serum antibody responses than IDV. However, by 12 DPI virus specific responses were higher in IDV infected animals in serum, BAL and nasal swabs compared to EAavH1N1.

### Cytokine responses following EAavH1N1 and IDV infection

Cytokine-secreting cells were quantified in BAL, tracheobronchial lymph nodes (TBLN), peripheral blood mononuclear cells (PBMC), and spleen using ELISpot IFNγ and IL-2 assays after *ex vivo* stimulation with live IDV or EAavH1N1 viruses **(Fig. 3A and S2)**. EAavH1N1 induced IFNγ ELISpot secreting cells in BAL and TBLN by 5 DPI which increased by 12 DPI and were significantly higher compared to IDV in BAL. In PBMC and spleen IFNγ ELISpot responses were comparable between IDV and EAavH1N1 at 5 and 12 DPI **(Fig. 3A)**. Interleukin 2 responses showed similar kinetics to IFNγ but were lower overall **(Fig. S2)**.

**Fig. 3:**
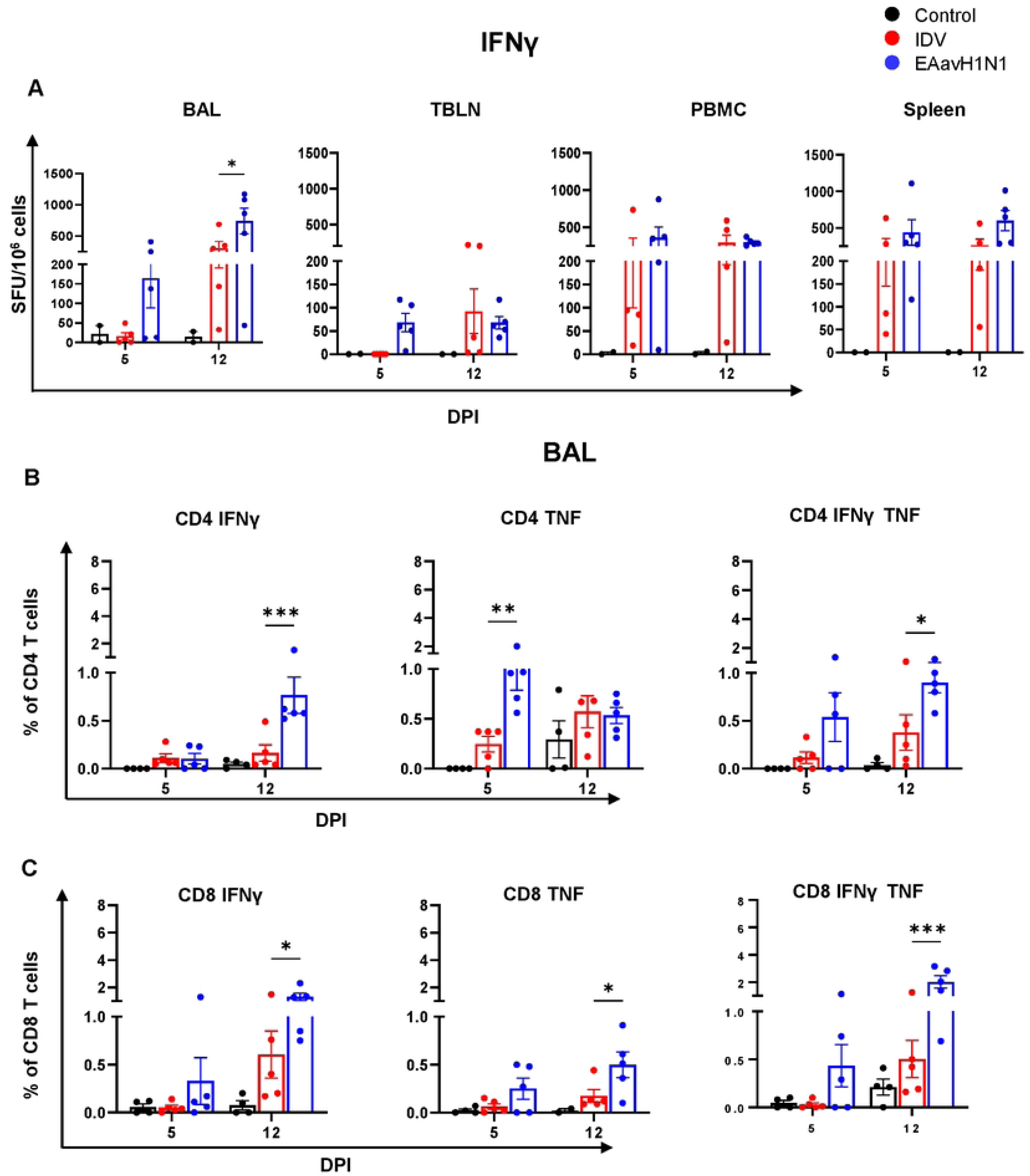
Virus-specific T-cell responses by ELISpot and Intracellular Cytokine Staining (ICS) : (A) Cells producing IFNy in BAL, TBLN, PBMC, and spleen in response to whole live IDV or EAavH1N1 virus at the indicated timepoints were measured by ELISpot. Symbols in A represent mean spot-forming units (SFU) per group, with error bars indicating. (B, C) IFNy- and TNF-producing co4• and cos• T cells were measured by intracellular cytokine staining following stimulation with whole virus at the indicated timepoints. Data are presented as mean ± SEM, with symbols indicating individual animals. Normally distributed data (A,-C) were analysed by two-way ANOVA followed by Bonferroni’s multiple comparisons test. Asterisks denote statistically significant differences between IDV and EAavH1N1 responses at indicated timepoint time points (**p < 0.05, ***p < 0.01, ****p < 0.001).

Intracellular cytokine staining (ICS) was performed to identify the responding IFNγ⁺ and TNF CD4⁺, CD8⁺, and γδ⁺ T cell subsets **(Figs. 3B, C and S3, S4, S5, S6**). In BAL, EAavH1N1 infection significantly increased the proportions of IFNγ⁺ and TNF⁺ CD4⁺ cells at 12 DPI compared to IDV and of IFNγ⁺TNF⁺ CD4⁺ T cells at 5DPI (**Fig. 3B**). CD8⁺ T cells exhibited a similar pattern, with significantly higher frequencies of IFNγ⁺, TNF⁺, and IFNγ⁺TNF⁺ CD8⁺ cells at 12 DPI in EAavH1N1-infected animals compared to IDV (**Fig. 3C**).

PBMC and spleen responses also showed a significantly higher proportions of cytokine secreting CD4⁺ and CD8⁺ cells in EAavH1N1 infected animals **(Fig. S4A-B)** Responses in TBLN were low, but still higher for both CD4 and CD8 cytokine producing cells in EAavH1N1 infected animals (**Fig. S5**). γδ T cell cytokine responses were low overall, but were higher in EA avian H1N1 infected pigs, especially in BAL and TBLN, except that IFNγ⁺TNF⁺ γδ⁺ T cells in IDV infected pigs in TBLN **(Fig. S6)**.

Overall, these data indicate that both viruses induced cytokine production in T cells following infection, with a higher proportion of IFNγ and TNF secreting cells observed in EA avian H1N1 infected animals compared to IDV.

### Differential abundance analysis reveals infection-specific cellular response in the porcine lung

Single cell (sc) RNA-seq was performed from lung tissue derived cell suspensions of control pigs (n = 3), IDV-infected pigs (n = 3 at 1 DPI; n = 4 at 12 DPI), and EAavH1N1-infected pigs (n = 4 at 12 DPI), resulting in 14 samples. EAavH1N1 samples at 1 DPI did not meet quality control standards and were excluded. After unsupervised clustering we identified three main compartments: myeloid, lymphoid, and non-immune cells with multiple cell subpopulations in each compartment (**Figs. 4A and S8**). Infection status significantly influenced composition of these three main compartments, accounting for 48% of total variance (PERMANOVA: F = 3.09, r² = 0.48, p = 0.022). IDV at 1 DPI was enriched for myeloid populations compared to control pigs, while both infections at 12 DPI showed increased lymphoid cell representation (**Fig. 4B**).

**Fig. 4:**
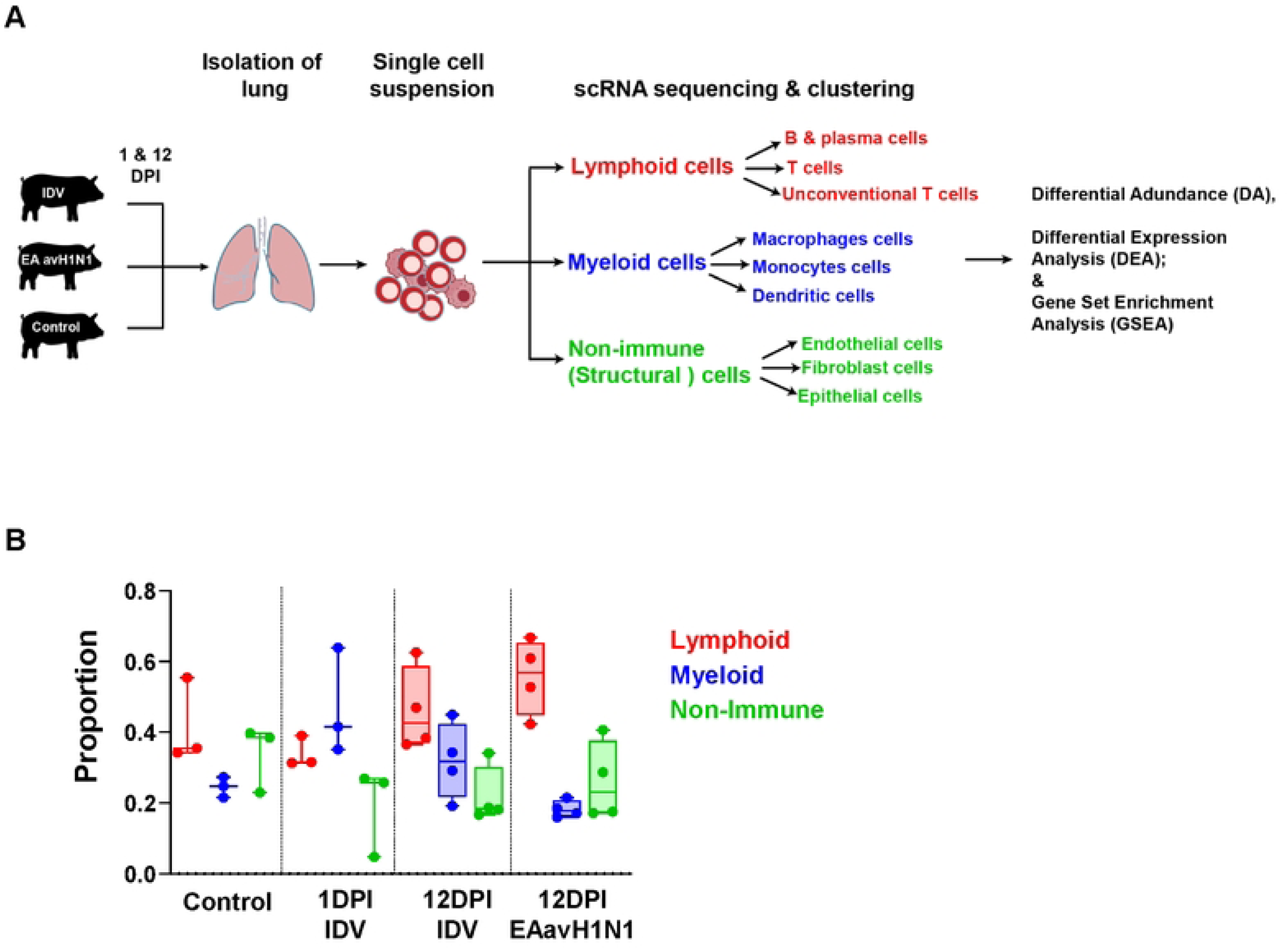
Single-cell transcriptomic profiling of porcine lungs following IDV and EAavH1N1 infection. (A) Schematic illustration of the experimental workflow. Pigs were intranasally infected with IDV or EAavH1N1, with uninfected pig lung serving as controls. Lungs were collected at 1 and 12 DPI and processed into single-cell suspensions for scRNA-seq. Unsupervised clustering identified major cellular compartments, including lymphoid cells, myeloid cells and non-immune structural cells. (B) Relative proportions of major cellular compartments in the lung across time point and infection group. Box plots show the distribution of lymphoid (blue), myeloid (red) and non-immune structural (grey) cells among total recovered cells per animal. Each dot represent individual animals.

### Reduced expression of immune genes and metabolic pathways in lung cell populations during IDV infection

Differential gene expression for IDV samples from 1 DPI compared to control samples showed reduced expression of genes across several cell types present within myeloid, lymphoid and non-immune cells (**Fig. 5**). Myeloid cells (**Fig. 5A, Table S1**) showed a modest but consistent reduced expression of genes linked to basic cellular functions, including pyrimidine metabolism (DPYS), stress-response chaperones (HSPA6), and lysosome-related vesicle trafficking (HPS5), suggesting reduced metabolic and stress-response activity (**Fig. 5B, Table S2**). In lymphoid populations (**Fig. 5C, Table S2**), a broader set of antiviral and interferon-associated genes (including DDX60, EIF2AK2, MX1, and RSAD2) was lower in IDV infected pigs, indicating a dampened antiviral response in early infection. A similar pattern was observed in non-immune cells, particularly in endothelial-like populations, which exhibited widespread reduced expression of interferon-stimulated and viral sensing genes involved in innate immune defence (e.g. ZBP1, IFIT1, IFIT2, IFIT3, DDX60, EIF2AK2, USP18, CMPK2, MX1, MX2, IFI44, IFI44L, IRF7, OAS1, OAS2, IFIH1, HERC5, HERC6, DHX58, RIGI/DDX58, ADAR, PLSCR1, PML and PARP9/PARP12/PARP14) (**Fig. 5D, Table S3**). This was also reflected in the gene set enrichment analysis (GSEA), where across the three compartments at least one cell population showed altered NOD-like receptor, RIG-I-like receptor and Toll-like receptor signalling or cytosolic DNA sensing (**Figs 5E, Table S4**). Overall, these findings suggest a coordinated suppression of antiviral and stress-response pathways across multiple lung cell compartments at this early stage of IDV infection.

**Fig. 5.**
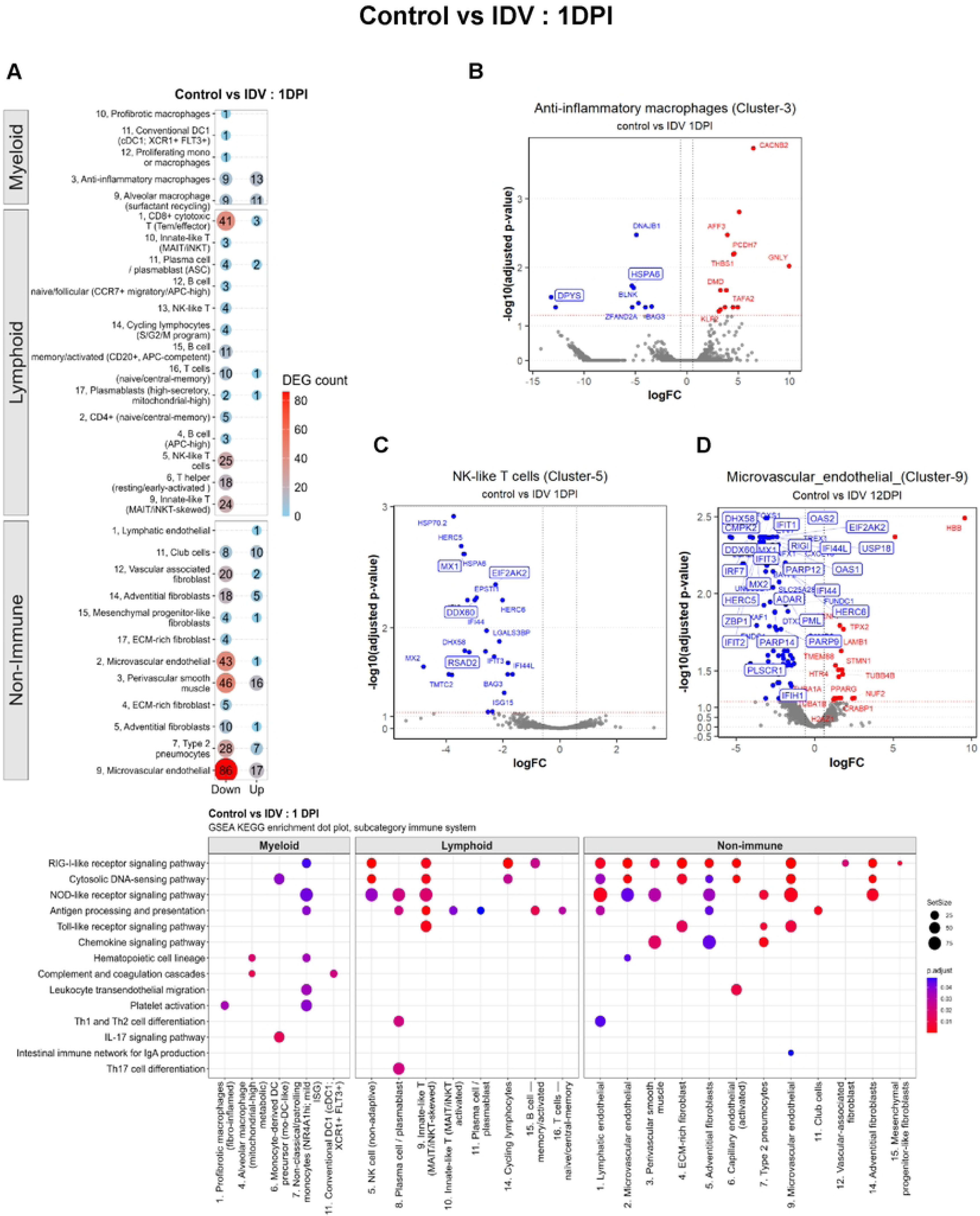
Myeloid, lymphoid, and non-immune cell programs show distinct responses during early IDV (1DPI) infection in pig lung. (A) Bubble heatmap shows the number of differentially expressed genes (DEGs) per subset compared to the control. Numbers preceding each cell type indicate the cluster number within each cell compartment. **(B-C)** Volcano plot displays a representative cell subtype, with each point representing a gene coloured by expression direction (red: upregulated; blue: downregulated; grey: not significant), plotted by log fold change (logFC) versus -log10 adjusted P value. (E) Dot plot shows KEGG GSEA results for the same cell compartment, where dot size indicates set size and dot colour reflects adjusted enrichment significance (p.adjust).

At 12 days post infection, some cell clusters across the three major cell compartment showed increased gene expression compared to control pigs, but reduced gene expression still dominated (**Fig. 6A**). Several myeloid cell clusters displayed a consistent downregulation of genes involved in transcriptional regulation (CEBPB) and core components of the translational machinery (RPL38, RPS29, SEC61G) (**Fig. 6B, Table S1**), indicating reduced protein synthesis and altered cellular activity. In lymphoid populations, a large set of shared DEGs across T, B, and NK cells was dominated by genes with reduced expression that are linked to mitochondrial respiration (ATP5ME, ATP5MK, COX17, COX6C, NDUFB1), ribosomal function (RPL37, RPL37A, RPL38, RPS25, RPS26, RPS28, RPS29), and immune signalling (CEBPB, JUNB, MAP2K3, ZFP36L2) (**Fig. 6C, Table S2**). This pattern suggests a broadly suppressed transcriptional and metabolic state across lymphoid lineages. Non-immune cells showed a similar trend (**Fig. 6A**), with reduced transcriptional expression of ribosomal (RPL38, RPS29, RPS28, RPL37, and RPL37A), protein-processing (SEC61G), and post-transcriptional regulatory genes (ZFP36L2) (**Fig. 6D, Table S3**). GSEA analysis showed similar pathways were affected as on 1DPI (e.g. NOD-like receptor signalling, Toll-like receptor signalling, cytosolic DNA sensing) across several cell clusters in the three cell compartments (**Fig. 6E, Tables S4, S5**) and additional chemokine signalling and antigen processing/ presentation. Overall, these findings point to a coordinated shift towards reduced protein synthesis and immune activity, combined with altered mitochondrial function, across multiple lung cell compartments at this later stage of infection.

**Fig. 6.**
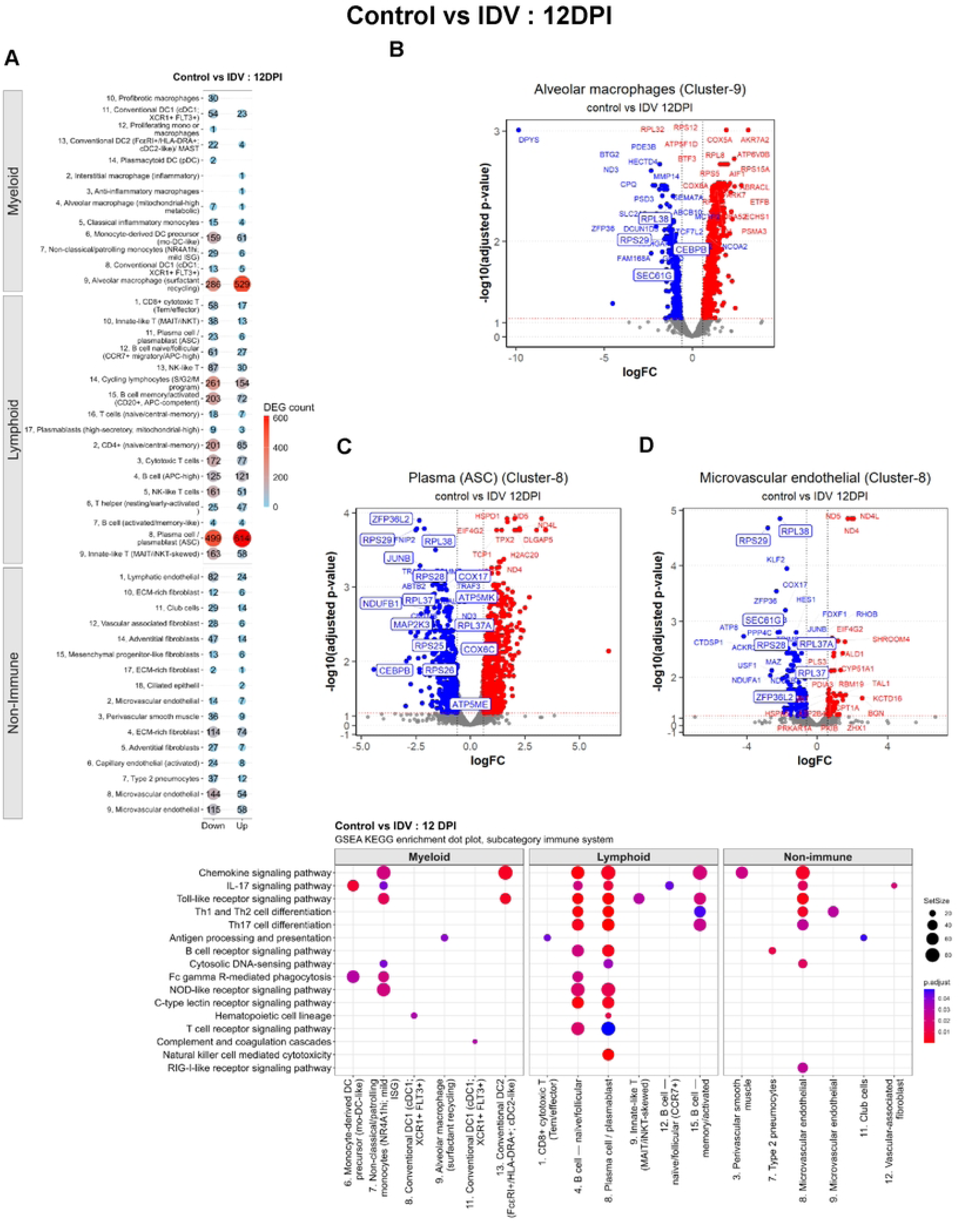
Myeloid, lymphoid, and non-immune cell programs show distinct responses during late IDV (12DPI) infection in pig lung. **(A)** Bubble heatmap shows the number of differentially expressed genes (DEGs) per subset in IDV infection group compared to the control. Numbers preceding each cell type indicate the cluster number within each cell compartment. **(B-C)** Volcano plot displays a representative cell subtype, with each point representing a gene coloured by expression direction (red: upregulated; blue: downregulated; grey: not significant), plotted by log fold change (logFC) versus -log10 adjusted P value. **(E)** Dot plot shows KEGG GSEA results for the same cell

### At 12DPI EA avian H1N1 infection is characterised by a recovery-associated gene pattern

Compared to IDV, the number of differentially expressed genes (DEGs) for EAavH1N1 infected pigs in relation to controls at 12 DPI was considerably lower across all three cell compartments (**Fig. 7A**). In myeloid cells, differential expression was primarily found in alveolar macrophage cluster 4, that showed an increased expression of NF-κB inhibitory and signalling modulators (NFKBIA, IL4R, MAPK6) alongside reduced expression of the pro-inflammatory transcription factor CEBPB (**Fig. 7B, Table S1**). In lymphoid populations, a reduced expression of genes involved in protein synthesis, metabolism, and immune signalling (MRPL44, PSMB3, ATP8, CEBPB) was found across T, B, and NK cell lineages, with only limited upregulation of genes linked to metabolic or chromatin-related functions (CYB5R3, H2AC20) (**Fig. 7C, Table S2**). Non-immune cells displayed a partially overlapping pattern, with reduced expression of metabolic and housekeeping genes (ATP8, SEC61G) alongside induction of stress- and signalling-related genes (JUN, ATF3, DUSP1) (**Fig. 7D, Table S3**), indicating tissue adaptation rather than active inflammation. While transcriptional data from EAavH1N1 infected pigs at 1 DPI could not be analysed, these findings suggest a transition from early antiviral responses to a more recovery- or resolution-associated state across lung cell compartments. This is also reflected in the GSEA results that shows an alteration of signalling pathways present in different cell types, ranging from innate immune cells to platelets and adaptive immune cells (**Fig. 7E, Tables S4, S5**).

**Fig. 7.**
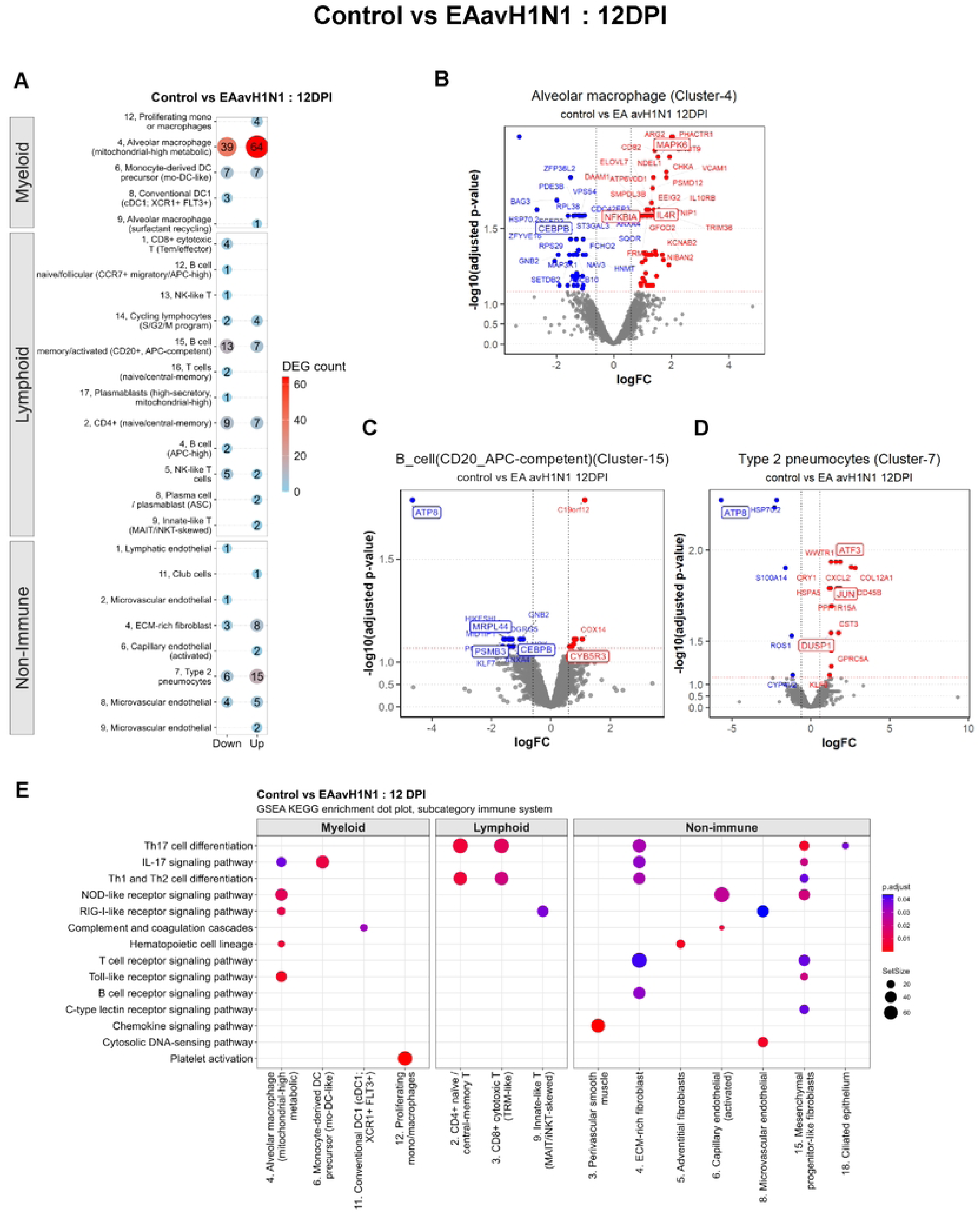
Myeloid, lymphoid, and non-immune cell responses during late EAavH1N1 (12DPI) infection in pig lung. (A) Bubble heatmap shows the number of differentially expressed genes (DEGs) per subset in EA avH1N1 infection group compared to the control. Numbers preceding each cell type indicate the cluster number within each cell compartment. **(B-C)** Volcano plot displays a representative cell subtype, with each point representing a gene coloured by expression direction (red: upregulated; blue: downregulated; grey: not significant), plotted by log fold change (logFC) versus -log10 adjusted P value. (E) Dot plot shows KEGG GSEA results for the same cell

### Differential metabolic and translational gene expression distinguishes IDV from EAavH1N1

Finally, we compared scRNA-seq data from 12 DPI from the two viral infections against each other, with EAavH1N1 as the baseline. The total number of DEGs was high across all the major cell types (**Fig. 8A**). In myeloid cells, the strongest changes were observed in interstitial macrophages, monocyte-derived DC precursors, and dendritic cell populations, showing increased expression of ribosomal and metabolic genes (e.g. RPS28, RPS29, ATP5ME, COX17) and reduced expression of mitochondrial-encoded genes (e.g. ND4, ND5) (**Fig. 8B, Table S1**). A similar pattern was found in lymphoid populations, where the highest DEG numbers were found in clusters representing plasmablasts, B cells, and CD8⁺ and CD4⁺ T-cell subsets, again characterised by increased expression of ribosomal and metabolic genes (e.g. RPS28, RPS29, ATP5ME) and reduced mitochondrial gene expression, with additional modulation of regulatory genes such as HES1 (**Fig. 8C, Table S2**).

**Fig. 8.**
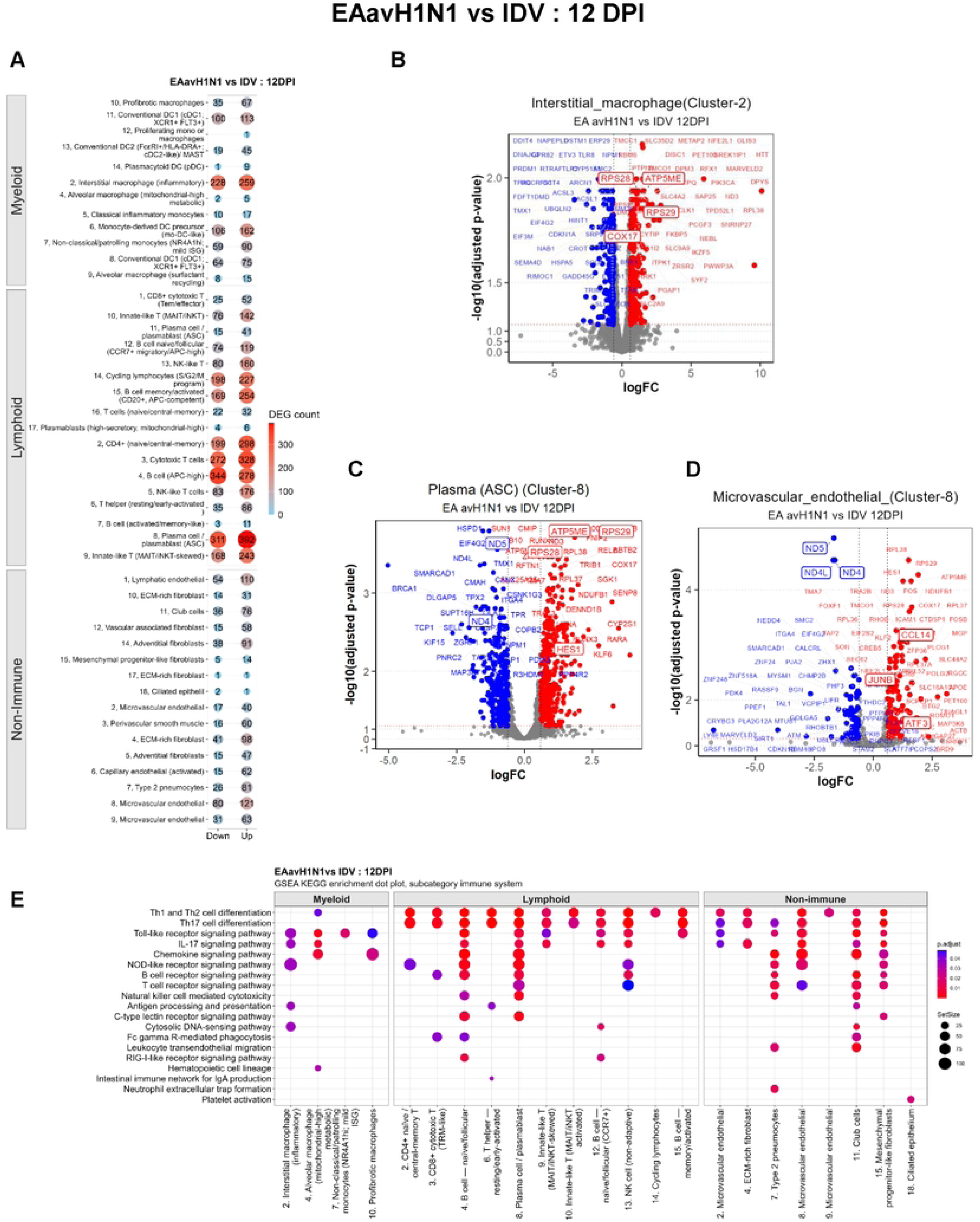
EAavH1N1 and IDV elicit distinct transcriptional and pathway responses in lung cell compartments at 12 DPI. **(A)** Bubble heatmap shows the number of differentially expressed genes (DEGs) per subset in IDV infection group compared to the EAavH1N1. Numbers preceding each cell type indicate the cluster number within each cell compartment. **(B-C)** Volcano plot displays a representative cell subtype, with each point representing a gene coloured by expression direction (red: upregulated; blue: downregulated; grey: not significant) in IDV infected group, plotted by log fold change (logFC) versus -10910 adjusted P value. **(E)** Dot plot shows KEGG GSEA results for the same cell compartment, where dot size indicates set size and dot colour reflects adjusted enrichment significance (p.adjust).

In non-immune cells, this pattern was most prominent in microvascular endothelial (cluster 8) and lymphatic endothelial cells (cluster 1), followed by fibroblast and perivascular populations, with increased expression of biosynthetic and signalling-associated genes (e.g. JUN, ATF3, CCL14) and reduced expression of mitochondrial and housekeeping genes (ND4, ND4L, ND5) (**Fig. 8D, Table S3**). This was also reflected in GSEA analyses, which showed that multiple signalling pathways present in innate and adaptive immune cells, but also other cell types were altered (**Fig. 8E, Tables S4, S5**). Overall, these findings indicate changes in cellular metabolism and protein synthesis across immune and structural cell types, distinguishing the IDV response strongly from EAavH1N1 at 12DPI.

## Discussion

The present study provides a comparative analysis of host responses to two antigenically and biologically distinct swine influenza viruses, showing clear differences in infection kinetics, respiratory pathology, and immune responses. The data indicates early, robust host responses associated with EAavH1N1 and a more prolonged, yet immunologically attenuated, profile following IDV infection.

IDV shedding persisted longer with higher titres, suggesting either more efficient replication or delayed clearance at mucosal surfaces. In contrast, EAavH1N1 triggered rapid tissue damage in both upper and lower respiratory tract compartments, indicating that early host responses contribute to pathology. These findings are broadly consistent with previous work in pigs and other animal models. Experimental infections with EAavH1N1 in swine show efficient replication in both upper and lower respiratory tract and the development of cranioventral bronchopneumonia [32, 33]. Studies of reassortant EAavH1N1 viruses in mice have further demonstrated that specific gene constellations, particularly those incorporating pandemic 2009 internal genes or polymerase mutations, significantly increase replication and inflammatory responses, leading to more severe lung pathology [9, 12, 34]. Ferret models have shown efficient replication and transmission with only moderate pathology, suggesting that host species and viral genotype together determine disease severity [35].

In contrast, the more prolonged but clinically attenuated profile for IDV observed here is in agreement with reports from cattle, where infection is typically mild, often restricted to the upper respiratory tract, and associated with efficient transmission but limited overt pathology unless compounded by coinfection [23, 27, 28]. Similarly in guinea pigs and ferrets, IDV has been shown to reach substantial viral loads in the respiratory tract, in some cases including the lower respiratory tract, yet with minimal clinical signs [36, 37].

The differences between IDV and EAavH1N1 are also reflected in the antibody responses. EAavH1N1 elicited earlier systemic IgG and IgA responses, as well as earlier neutralizing activity, consistent with faster antigen presentation and immune priming. In contrast, IDV induced a delayed but ultimately higher antibody response in serum, BAL and nasal swabs at 12 DPI. This likely reflects prolonged antigen exposure due to sustained viral replication. The absence of detectable neutralizing activity in mucosal samples, despite measurable IgA, suggests that local antibodies may be non-neutralizing, below detection thresholds because of the diluted nature of the samples.

The two infections were further distinguished by their cellular immune responses. EAavH1N1 induced higher numbers of IFNγ and TNF producing cells in BAL and TBLN, with responses also detected in PBMC and spleen, indicating a stronger activation of adaptive immunity. In contrast, IDV elicited weaker cytokine responses despite prolonged viral presence. The strong IFNγ and TNF responses induced by EAavH1N1 in this study are consistent with observations in more pathogenic influenza A infections, including murine models of reassortant EAavH1N1 where elevated IL-6, TNF, IFNγ and chemokine responses correlate with increased lung pathology [9, 12].

Single cell transcriptomic analysis provided mechanistic insights into these observations. As early as 1 day post infection, IDV infection was associated with widespread downregulation of interferon stimulated genes and antiviral pathways across myeloid, lymphoid, and structural cell populations. Key components of viral sensing and innate immune activation, including RIG I like receptor and Toll like receptor signalling pathways, were reduced. While correlative, these findings suggest that IDV can dampen early innate immune responses, which may contribute to higher viral load and prolonged shedding. At the same time, this reduced antiviral signalling may limit inflammation and contribute to the milder lung pathology observed in IDV infected pigs. The early downregulation of interferon stimulated genes and innate sensing pathways during IDV infection is consistent with experimental evidence in cattle indicating limited type I interferon activation and with the known role of viral proteins in suppressing host responses [27, 29, 38].

This dampened transcriptional activity by IDV persisted at 12 DPI, extending to genes involved in protein synthesis, mitochondrial function, and immune signalling across all major lung cell compartments. Such coordinated downregulation indicates a sustained alteration of lung tissue homeostasis and may explain the reduced local T cell responses in BAL, potentially reflecting a microenvironment less supportive of type 1 T cell response. Despite this, IDV infected pigs showed higher antibody titers in serum and BAL. One possible explanation is that prolonged viral replication enhances antigen delivery to draining lymph nodes, this may have supported B cell activation and antibody production, even if local lung responses were dampened.

In contrast, EAavH1N1 infection at 12 DPI showed fewer transcriptional changes relative to controls, with gene expression patterns consistent with resolution and tissue repair. Reduced expression of pro-inflammatory and metabolic genes, alongside induction of regulatory and stress response pathways, suggests that viral clearance had occurred and that lung tissue was transitioning towards recovery.

Overall, these findings support a model in which EAavH1N1 induces rapid and strong immune activation that drives early respiratory pathology but enables efficient viral clearance and recovery. In contrast, IDV causes a more prolonged infection characterised by higher viral replication, delayed and weaker immune responses, and sustained suppression of antiviral and metabolic pathways. These differences have important implications for transmission dynamics, disease outcome, and immune control of influenza viruses in swine and are also relevant for human health in settings with frequent pig-human interactions, such as in pig farming.

## Materials and Methods

### Viruses

This study used IDV (D/swine/Netherlands/PS-497/2021, GenBank accession numbers OP474071–OP474077) and EAavH1N1 (A/swine/Gent/274/2020, GenBank accession numbers OP458671–OP458678) [6]. Both viruses were propagated and quantified in Madin-Darby Canine Kidney (MDCK) cells. Viral titers were determined by plaque assay and reported as plaque-forming units per millilitre (PFU/mL).

### IDV and EAavH1N1 infection

All experiments were approved by The Pirbright Institute Ethical Review Committee and conducted in accordance with the U.K. Government Animal (Scientific Procedures) Act 1986 under Project License PP2064443. In all experiments, animals were acclimatized for at least 7 days and randomized into different groups and pens using Excel by the animal services staff. Researchers processing the samples were only aware of the pig numbers, not the group assignments. The pathologists were blinded to the group allocation when assessing the samples for gross pathology during *post-mortem* examination and subsequently during the histopathological assessment. All conditions were identical between the groups.

For infection studies, 36 five-week-old female 50% Landrace × 50% White Durocs cross (PIC Camborough 50) pigs (9–13 kg; mean 11.8 kg) were sourced from <u>Home - PIC UK</u> high-health commercial herd. All animals were seronegative for influenza A and D virus antibodies by ELISA with H1N1pdm09, IDV, and EAavH1N1. Pigs were randomly assigned to three groups: IDV-infected (n = 15), EAavH1N1-infected (n = 15) and untreated controls (n = 6). Pigs were intranasally infected with 6 × 10⁶ PFU/mL of either IDV or EAavH1N1 or DMEM w/o antibiotics, 1.5 mL per nostril, using a mucosal atomization device (MAD; Wolfe-Tory Medical) following sedation following sedation with a 3 mg/kg Zoletil and 1.5 mg/kg Stresnil. Clinical parameters including demeanour, appetite, respiratory signs, sneezing, coughing, nasal and eye discharge, faeces consistency, and rectal temperature were assessed. No significant clinical signs were observed in any of the pigs following the viral inoculation.

Nasal swabs were collected daily for 10 days post-infection (DPI) to assess viral load by plaque assay and RT-qPCR. Pigs were humanely euthanized with intravenous pentobarbital (Dolethal, 200 mg/mL) on days 1, 5, and 12 DPI, along with uninfected controls. At post-mortem, serum, PBMCs, lungs, tracheobronchial lymph nodes (TBLN), spleen, and bronchoalveolar lavage (BAL) were collected for analysis of viral load, pathology, immune responses and scRNA seq analysis.

### Lung gross pathology, histopathology, and immunohistochemistry

Gross and histopathological analyses were performed as previously described [39]. Lungs were removed, and dorsal and ventral aspects were evaluated and digital images acquired to quantify the surface area showing pneumonia using image analysis software (Nikon NIS-Ar). Samples from the cranial, middle, and caudal lobes of the right lung were collected into 10% neutral-buffered formalin, paraffin-embedded, sectioned at 4 μm, and stained with haematoxylin and eosin (H&E). Immunohistochemistry (IHC) for IDV and EAavH1N1 viruses’ nucleoprotein (NP) was performed on parallel sections [39, 40]. Histopathological changes were evaluated by a veterinary pathologist blinded to treatment groups. Lung histopathology was scored across five parameters: bronchiolar epithelial necrosis, airway inflammation, perivascular/bronchiolar cuffing, alveolar exudates, and septal inflammation on a 0 -4 scale, yielding a maximum of 20 per lobe (∼1.5 × 3 × 1.5 cm²) and 60 per animal. A cumulative score from the three lobes was calculated for each animal. Viral antigen distribution was additionally evaluated using the “Iowa” scoring system. IHC labelling was assessed separately in bronchioles and alveoli, with each scored 0–3 [41]. The same scoring system was used to evaluate lesions and presence of virus NP in the respiratory and olfactory nasal epithelium.

### Sample processing and cell isolation

Serum, nasal swabs, PBMC, BAL were collected and processed as previously described [39, 42]. Mononuclear cells were isolated from lungs, TBLN, and spleens. TBLN were minced, filtered through 70- and 40-μm strainers, washed, and resuspended in complete RPMI with 2% fetal bovine serum (FBS) and 1X Penicillin-Streptomycin (PS) [42]. Spleens were mechanically dissociated, filtered, centrifuged at 500 × g for 10 minutes, and mononuclear cells enriched using a Histopaque-1077 density gradient. Red blood cells (RBCs) were lysed with ammonium chloride lysis [43]. Lung-derived cells from medial and diaphragmatic lobes were digested with collagenase (0.7 mg/mL) and DNase (0.03 mg/mL), filtered, and RBCs lysed before resuspension in DMEM containing 10% FBS, 1% PS, and 1% L-glutamine. All cells were cryopreserved in FBS with 10% DMSO and stored in liquid nitrogen until use.

### Plaque assays

Viral load in nasal swabs was measured by plaque assay using MDCK cells. Samples were serially diluted in DMEM with 0.1% BSA and inoculated onto confluent MDCK monolayers. After 1 hour at 37 °C with 5% CO₂, the inoculum was removed, cells were washed with Dulbecco’s Phosphate-Buffered Saline (DPBS), and an overlay medium containing DMEM, 0.3% BSA, 2.86 mM L-glutamine, 0.3% sodium bicarbonate, 14 mM HEPES, 0.007% dextran, 1X PS, and 0.6% agar (Oxoid) was added. Plates were incubated for 72 h at 37 °C with 5% CO₂. Plaques for EAavH1N1 were visualized using was visualized using 0.1% crystal violet [39]. For IDV, plaques were detected by immunoperoxidase staining due to the lack of distinct cytopathic plaques [17, 44]. Briefly, cells were fixed with 500 μL of 30% formalin (F8775, Sigma Aldrich) in PBS at 4 °C overnight, washed, and permeabilized with 0.1% Triton X-100 (T9284, Sigma Aldrich) in PBS for 15 minutes at room temperature. After blocking with SuperBlock™ Blocking Buffer (37515, Thermo Fisher Scientific), cells were incubated with cattle anti-IDV immune serum (kindly provided by Dr. Mariette Ducatez, University of Toulouse, France) for 60 minutes, washed, and then incubated with HRP-conjugated goat anti-cattle IgG-Fc (A100-104P, Bethyl Laboratories, Inc.) for 60 minutes. Plaques were visualized using 3-amino-9-ethylcarbazole (AEC) chromogenic substrate (AEC101, Sigma-Aldrich) according to the manufacturer’s instructions. Viral titer were calculated as log₁₀ PFU/mL by averaging plaque counts and adjusting for dilution.

### RNA isolation and quantification of viral genomic RNA

Genomic RNA (gRNA) copies of IDV-NP and EAavH1N1-M1 were quantified in NS by qRT-PCR [43]. Viral RNA was extracted from 200 μL of each sample and from purified virus stocks using the QIAamp Viral RNA Mini Kit (52906, Qiagen) according to the manufacturer’s instructions. Standards were generated by isolating gRNA from purified IDV and EAavH1N1 viruses. Copy numbers per μL were calculated, and 10-fold serial dilutions from 1 × 10¹⁰ to 10 copies were prepared. Standard dilutions were included on each qRT-PCR plate with test samples. Reverse transcription and amplification were performed using the EXPRESS One-Step Superscript™ qRT-PCR Kit (11781200, Invitrogen) on an AriaMx Real-time PCR System (Agilent), following the thermal cycling conditions in **Table S6.** Custom gene expression assays were designed using sequences from GenBank, NCBI (accession numbers OP474075.1, OP458677.1, MW362720.1, OP458613.1, OP458637.1, OP458685.1, MW362640.1). Primer and probe sequences are listed in **Table S6**. Each sample and standard was run in duplicate. Standard curves were generated for each plate, and PCR efficiency (E) was calculated from the slope of the regression line using E = 10(−1/slope). Viral RNA copy numbers in experimantal samples were determined by extrapolating cycle threshold (Ct) values against the standard curve and expressed as log₁₀ NP or M1 gRNA copies per mL of sample. No-template controls (NTCs) were included on each plate. All experiments were conducted in compliance with MIQE (Minimum Information for Publication of Quantitative Real-Time PCR Experiments) guidelines [45, 46].

### Enzyme-linked immunosorbent assays (ELISA)

Endpoint ELISAs were used to quantify IDV- and EAavH1N1-specific IgG and IgA antibodies in serum, BAL, and nasal swabs, as previously described [31, 35, 47]. Serum was collected on days 1, 5, 7, and 12 DPI. BAL on days 1, 5, and 12 DPI; and NS on days 1, 5, 7, and 10 DPI. Ninety-six–well plates were coated overnight at 4 °C with IDV or EAavH1N1 virus in 0.05 M carbonate - bicarbonate buffer (pH 9.6) (Sigma Aldrich). Plates were washed with DPBS containing 0.05% Tween-20 (Sigma Aldrich) and blocked with 4% skimmed milk in PBS w/ 0.05% Tween-20 for two hours at room temperature. Two-fold serial dilutions of heat-inactivated serum (starting at 1:20), BAL (1:2), or nasal swabs (1:8) were prepared in blocking buffer and incubated in duplicate (one hour at room temperature for serum and BALF; 16 hours for nasal swabs). Negative controls were included on each plate. After washing, plates were incubated with mouse anti-pig IgG-HRP (1:20,000) or anti-pig IgA-HRP (1:10,000) (A100-102P, Bethyl Laboratories, Inc) in blocking buffer for one hour at room temperature, washed, and developed with TMB substrate (50 µL/well) (34029, Thermo Scientific). Reactions were stopped with 1M H₂SO₄ (50 µL/well), and absorbance was measured at 450 nm with a 630 nm reference using a VersaMax microplate reader (Molecular Devices). Endpoint titers were defined as the log₁₀ of the highest dilution exceeding the cut-off (mean plus three standard deviations of negative controls or control animals) [48].

### Microneutralization assay

Neutralising antibody (nAb) titers in serum and BAL were measured using a microneutralization (MN) assay as previously described [43, 46, 49]. Samples were heat-inactivated at 56 °C for 30 minutes, serially diluted two-fold in virus growth medium (VGM; DMEM with 0.2% BSA, 1× PS, and 2 μg/mL TPCK-trypsin), and combined with live IDV or EAavH1N1 viruses. The initial dilution was 1:20 for serum and 1:2 for BAL. Viruses were diluted to 1:50 (IDV) or 1:200 (EAavH1N1) in VGM. Control wells included medium only (negative) and virus + cells only (positive). After a 2-hour incubation at 37 °C with 5% CO₂, 3 × 10⁴ MDCK cells per well were added in VGM, and plates were incubated overnight under the same conditions. Following incubation, cells were fixed with cold 4% paraformaldehyde (J61984.AP, Thermo Scientific) for 15 minutes, permeabilised with DPBS containing 0.05% Triton X-100 and 0.75 M glycine (50046, Sigma-Aldrich) for 30 minutes at room temperature, and washed with DPBS-T. Plates were incubated with primary antibodies [anti-influenza A NP (1:3500) (MCA2751, Bio-Rad) and cattle anti-IDV serum (1:200)] in SuperBlock™ buffer, followed by HRP-conjugated anti-mouse IgG (1:3500) (STAR207P, Bio-Rad) or anti-cattle IgG (MCA2439P, Bio-Rad) (1:1000) in SuperBlock™ buffer for 1 hour at room temperature. After washing, plates were developed with TMB substrate (50 μL per well), and reactions were stopped with 1 M H₂SO₄ (50 μL per well). Absorbance was measured at 450 nm with a 630 nm reference using a VersaMax microplate reader (Molecular Devices). MN titers were defined as the log10 50% inhibitory dilution (log10ID₅₀), calculated as the dilution corresponding to the midpoint between the optical densities of virus-infected (positive control) and uninfected (negative control) wells, determined by linear interpolation from the titration curve.

### Enzyme-linked immunospot (ELISpot)

Cryopreserved cells from BAL, TBLN, spleen, and PBMC were used to assess IFNγ- and IL-2 producing cell frequencies by ELISpot, as previously described [31, 47]. Multiscreen® 96-well plates (MCE membrane, MAHAS4510; Merck Millipore) were coated overnight at 4 °C with either 1 μg/mL anti-pig IFNγ (clone P2G10; BD Pharmingen) in carbonate–bicarbonate (CBC) buffer or 10 μg/mL anti-pig IL-2 (clones MT264/267; Mabtech) in DPBS. Plates were washed with PBS and blocked with complete culture medium (RPMI 1640 with 10% FBS, 1X PS) for 2 h at 37 °C. Cells (2.5 × 10⁵/well) were seeded in triplicate and stimulated with live IDV or EAavH1N1 (MOI = 1), Concanavalin A as a positive control (00-4978-03, Invitrogen), or culture medium as a negative control. After 48 h incubation at 37 °C with 5% CO₂, plates were washed with PBS-T and incubated for one hour at room temperature with biotinylated anti-pig IFN-γ (0.5 μg/mL; clone P2C11, BD Pharmingen) or anti-pig IL-2 (0.5 μg/mL; clone MT265, Mabtech). Plates were washed and incubated with streptavidin–alkaline phosphatase (1:1000; S291, Invitrogen) for one hours at room temperature, then developed with BCIP/NBT substrate (1706432, Bio-Rad) according to the manufacturer’s instructions. Reactions were stopped after 20 min with tap water, and plates were air-dried. Spots were counted using the CTL ImmunoSpot® M6 automated analyser (CTL, Cleveland, OH, USA) and analysed with ImmunoSpot® Software version 7.1. Results were presented as the number of IFNγ– or IL-2–producing cells per 10⁶ stimulated cells, after subtracting background spots from medium controls.

### Intracellular cytokine staining (ICS)

IDV and EAavH1N1-specific CD4⁺, CD8⁺, and γδ T-cell responses were quantified by ICS assays using cells isolated from BAL, TBLN, PBMC, spleen, and MLN as previously described [31, 47]. For each tissue, 2 × 10⁶ cells were resuspended in 200 μL RPMI with 10% FBS and 1× PS, then exposed to one of three conditions: (i) mock unstimulated, (ii) virus-stimulated (MOI = 1), or (iii) PMA + ionomycin (1×; 00497593, Invitrogen) as a positive control. Virus-stimulated and mock cultures were incubated at 37 °C for 16 h, while PMA + ionomycin cultures were incubated for four hours in a CO₂ incubator. GolgiPlug (555029, BD Biosciences) was added four hours before staining. After stimulation, cells were washed in DPBS or BD Perm/Wash buffer (555028, BD Biosciences) as appropriate. All staining steps were performed in 50 μL with pre-titrated antibodies. Cells were harvested (520 x g, 5 min, 4 °C), washed twice in DPBS, and stained with mouse anti-γδ TCR for 20 min at 4 °C. Following another wash, cells were incubated with anti-mouse IgG–BUV395 for 20 min at 4 °C in the dark, then blocked with anti-mouse serum and washed twice with DPBS. Surface staining was performed with LIVE/DEAD™ Fixable Near-IR (Invitrogen), CD3, CD4α, and CD8β for 20 min at 4 °C in the dark. After two additional DPBS washes, cells were fixed and permeabilized with BD CytoFix/CytoPerm for 15 min at 4 °C. Intracellular staining for IFN-γ and TNF was conducted in 1× Perm/Wash buffer for 30 min at 4 °C in the dark. Cells were washed twice, resuspended in 200 μL 1× Perm/Wash buffer. Further details of antibodies used are listed in Table S6. Samples were acquired on a Cytek Aurora spectral flow cytometer, unmixed by SpectroFlo software (Cytek) using single-stained controls, and analyzed in FlowJo v10 (BD Biosciences).

### Single cell RNA-Seq library preparation and sequencing

Single-cell suspensions were prepared from lung tissues of twenty pigs for transcriptomic profiling using the 10x Genomics 3′ Gene Expression platform. Suspensions were adjusted to 10,000 cells per sample and encapsulated with the Chromium iX Controller, Next GEM Chip G, and the Single Cell 3′ v3.1 library preparation kit (10x Genomics, Pleasanton, CA, USA). Library quality and concentration were evaluated using the Qubit DNA Broad Range assay (Life Technologies, UK), Illumina Library Quantification Kit (New England Biolabs, USA), and TapeStation D5000 DNA assay (Agilent Technologies, USA). Libraries were pooled and sequenced on two NextSeq 2000 P4 100-cycle runs (Illumina, San Diego, CA, USA), each with 1% PhiX control, targeting approximately 200 million reads per sample. Sequencing was performed according to the manufacturer’s recommendations.

### Single-cell RNA-Seq data pre-processing and quality control

Single-cell RNA-seq profiles from twenty samples were analysed. Raw BCL files were demultiplexed using the mkfastq pipeline in Cell Ranger v7.1.0 (10x Genomics). Reads were aligned to the Sus scrofa reference genome (assembly 11.1; GCA_000003025.6) using STAR v2.7.2a within Cell Ranger, and UMI counts were generated with the Cell Ranger count function. All downstream analyses were conducted in R v4.3.2. Stringent filtering was applied to ensure data quality and minimise technical artefacts. Empty and barcode-swapped droplets were removed using DropletUtils v1.22.0 [50, 51]. Cells with total UMI counts above 50,000 or below 1,000 were excluded to remove potential multiplets and low-quality cells. Additional filtering excluded cells with fewer than 500 detected genes, high mitochondrial gene content (>20%), or those predicted as doublets by scds v1.18.0. Genes with zero expression across all cells were also excluded. Library size differences were corrected using scaling normalisation with computePooledFactors from the scuttle package v1.12.0, followed by log2 transformation with a pseudo-count of 1 [52]. Fourteen samples from three independent sequencing runs were integrated using the fastMNN algorithm in the batchelor package v1.18.1 to correct for batch effects [53].

### Clustering and annotation of scRNA-seq libraries

Shared nearest-neighbour (SNN) graph-based clustering was performed using the clusterCells function in bluster v1.12.0. The top 5,000 highly variable genes were identified with getTopHVGs from the scran package v1.30.2 [54]. Clustering was based on a Jaccard-weighted graph and the Louvain community detection algorithm, with k set to 25 nearest neighbours. This process identified 37 transcriptionally distinct clusters. Clusters were annotated using canonical marker genes and grouped into three main cell compartments: myeloid, lymphoid (T, B, and NK-like T cells), and non-immune structural cells. Differences in immune cell composition between groups were assessed using permutational multivariate analysis of variance (PERMANOVA) in the vegan package v2.7-2. Prior to analysis, cell proportion data were transformed using centred log-ratio (CLR) scaling with the compositions package v2.0-9. Data visualisation and statistical summaries were generated using dplyr v1.1.4, tidyr v1.3.1, ggplot2 v3.4.3, and rstatix v0.7.3. Within each subset, cells were re-integrated and re-clustered using the same analytical framework to improve resolution. In the myeloid subset, 18 refined clusters were identified and annotated. Clusters 21 and 22, considered probable contaminants, were excluded from the final UMAP visualisation. In the lymphoid subset, 19 clusters were identified, representing activated, cytotoxic, memory, and unconventional T-cell populations. In the non-immune structural compartment, 20 clusters were resolved. Cluster 13, identified as immune cell-derived noise, was removed from the final UMAP representation because it did not correspond to structural cell types. Across all subsets, clusters were manually annotated using conventional cell-type nomenclature based on marker gene expression (Table-S6). For each cluster, the most informative differentially expressed genes were identified using the scoreMarkers function in scran v1.30.2 to define cluster-specific transcriptional signatures.

### Cell type scRNA-seq differential abundance and expression analyses

Differential abundance (DA) of cell subpopulations was evaluated across three major cell compartments under each experimental condition (control, EAavH1N1-, and IDV-infection) at 1 and 12 DPI. DA testing used edgeR v4.6.3 in R, applying a negative binomial generalised linear model (GLM) with empirical Bayes quasi-likelihood F-tests. The model included experimental condition as the main factor and sequencing run as a covariate to address batch effects. Sequencing depth was adjusted by including an offset based on cluster-level cell counts scaled to the total number of cells per sample. To prevent overfitting in clusters with limited replication, trend estimation was disabled during dispersion and quasi-likelihood dispersion estimation. Multiple testing was corrected using the Benjamini–Hochberg method. Pairwise DA comparisons included: at 1 DPI, IDV-infected (n = 3) vs control (n = 3); at 12 DPI, IDV-infected (n = 4) vs control (n = 3), EAavH1N1-infected (n = 4) vs control (n = 3), and IDV-infected (n = 4) vs EAavH1N1-infected (n = 4). Temporal changes within the IDV group were assessed by comparing samples at 1 DPI (n = 3) and 12 DPI (n = 4). Differential expression (DE) within each cell cluster was analysed using a pseudo-bulk approach, aggregating raw counts from all constituent cells to generate cluster-level profiles. Cluster–sample combinations with fewer than ten cells were excluded. Negative binomial GLMs were fitted for each cluster, and DE between experimental groups was assessed using empirical Bayes quasi-likelihood F-tests in edgeR v4.6.3, with sequencing run as a covariate. Multiple testing correction used the Benjamini–Hochberg procedure. Complete DE results are summarized in Tables S1, S2, and S3.

### Gene function and pathway enrichment scRNA-seq analysis

Adjusted p-values and log₂ fold-change estimates from edgeR analyses were used for downstream functional and pathway enrichment analyses with clusterProfiler v4.15.1. Gene set enrichment analysis (GSEA) was performed using KEGG pathway annotations [55, 56], applying a permutation-based approach to ranked gene lists ordered by log₂ fold change to identify pathways with coordinated transcriptional changes. Resulting p-values were corrected for multiple testing using the Benjamini–Hochberg method. Significantly enriched KEGG pathways (q < 0.05) were visualised using bubble plots. Complete GSEA results are provided in Table S4 and S5.

## Statistical Analysis

Statistical analyses were performed using GraphPad Prism v10.5.0 (GraphPad Software, San Diego, CA, USA). Data normality was assessed prior to analysis. Parametric data were evaluated with unpaired t-tests, one-way ANOVA, or two-way ANOVA, followed by Tukey’s or Dunnett’s post hoc tests. Non-parametric data were analysed using the Kruskal–Wallis test with Dunn’s multiple comparisons test. Statistical significance was set at (*p < 0.05, **p < 0.01, ***p < 0.001, ****p < 0.0001), and the specific test used is indicated in the figure legends.

## Conflict of interests

All authors declare no conflict of interests.

## Acknowledgments

We thank the animal staff at The Pirbright Institute for their exemplary animal care and acknowledge the support of the Bioinformatics Science Technology Platform. This research was supported by the UKRI Biotechnology and Biological Sciences Research Council (BBSRC) IAA award BB/X019780/1 as part of the EPICVIR project at the International Collaboration of Research on Infectious Animal Diseases (ICRAD). Research at Pirbright received funding from BBSRC through the Pirbright Institute’s Strategic Programme Grants (ISPGs) [BBS/E/PI/230002A; BBS/E/PI/230002B and BBS/E/PI/230002C], as well as the BBSRC National Bioscience Research Infrastructure: High Containment and Low Containment Services and Science Platforms grants [BBS/E/PI/23NB0004, BBS/E/PI/23NB0003]. We thank Dr. Mariette Ducatez from the University of Toulouse, France, for providing the anti-IDV cattle serum.

## Data availability

All R scripts used for processing and analysis of the scRNA-seq data are available at: XXXXXXXXX. Fastq and h5 files from sequencing are available at:GEO (GSEXXXX) .

## Author Contributions

**Conceptualization:** Elma Tchilian

**Data curation:** Ashutosh Vats, Liu Yang, Graham Freimanis, Francisco J. Salguero.

**Formal analysis:** Ashutosh Vats, Liu Yang, Francisco J. Salguero.

**Funding acquisition:** Kristien van Reeth, Elma Tchilian

**Investigation:** Ashutosh Vats, Liu Yang, Ehsan Sedaghat-Rostami, Tim Downing, Kristien van Reeth, Francisco J. Salguero, Wilhelm Gerner, Elma Tchilian

**Methodology:** Ashutosh Vats, Liu Yang, Ehsan Sedaghat-Rostami, Basudev Paudyal, Emily Briggs, Catherine Hatton, Emily Briggs, Graham Freimanis, Tim Downing, Kristien van Reeth, Francisco J. Salguero, Wilhelm Gerner, Elma Tchilian

**Project administration:** Elma Tchilian

**Resources:** Kristien van Reeth, Elma Tchilian

**Software:** Ashutosh Vats, Liu Yang, Francisco J. Salguero

**Supervision:** Ashutosh Vats, Basudev Paudyal, Wilhelm Gerner, Elma Tchilian

**Validation:** Ashutosh Vats, Liu Yang, Francisco J. Salguero, Wilhelm Gerner, Elma Tchilian

**Visualization:** Ashutosh Vats, Ehsan Sedaghat-Rostami, Liu Yang, Francisco J. Salguero, Wilhelm Gerner, Elma Tchilian

**Writing – original draft:** Ashutosh Vats, Wilhelm Gerner, Elma Tchilian

**Writing – review & editing:** Ashutosh Vats, Ehsan Sedaghat-Rostami, Liu Yang, Catherine Hatton, Emily Briggs, Graham Freimanis, Tim Downing, Kristien van Reeth, Francisco J. Salguero, Wilhelm Gerner, Basudev Paudyal, Elma Tchilian

## Notes

### Competing Interest Statement

The authors have declared no competing interest.

